# AUTO-SP: automated sample preparation for analyzing proteins and protein modifications

**DOI:** 10.1101/2025.01.08.631960

**Authors:** T. Mamie Lih, Liyuan Jiao, Lijun Chen, Jongmin Woo, Yuefan Wang, Hui Zhang

## Abstract

Liquid chromatography (LC) tandem mass spectrometry (MS/MS) is one of the widely used proteomic techniques to study the alterations occurred at protein expression level as well as post-translation modifications (PTMs) of proteins that are relevant to different physiological or pathological statuses. The mass spectrometric analysis of peptides fragmented from proteins (bottom-up proteomics) has emerged as one of the major approaches for proteomics. In this approach, proteins are first cleaved into peptides for mass spectrometric analysis and peptides with PTMs are further enriched followed by the LC-MS/MS analysis. To achieve a reproducible and quantitative proteomic characterization, a well-established protease digestion and PTM peptide enrichment protocol is critical. In this study, we developed an automated sample preparation (AUTO-SP) for analyzing proteins and protein modifications utilizing Patient-Derived Xenograft (PDX) breast cancer tumors (basal-like and luminal subtypes). The protein amount was quantified and proteins were further digested by using AUTO-SP for each PDX sample. Based on the data-independent acquisition (DIA)-MS data, we observed samples of the same breast cancer subtypes were highly correlated (>0.98). Additionally, >14,000 ubiquitinated peptides were identified in the PDX samples when using AUTO-SP for ubiquitin enrichment, while unique pathways were enriched from the basal-like and luminal subtypes. AUTO-SP demonstrated its efficacy to provide reliable and reproducible sample preparation procedure for MS-based proteomic and PTM analyses.

## INTRODUCTION

Quantitative proteomic analysis via mass spectrometry (MS) is one of the widely adapted techniques to study the alterations occurred at protein level as well as aberrant post-translation modifications (PTMs) of proteins that are relevant to the development of diseases^1-5^. Moreover, large-scale proteomic studies of several cancer types to understand the molecular basis of cancers have been feasible because of the recent advancements of MS^1-3,6-25^. To successfully conducting MS-based quantitative proteomic and PTM studies, the samples are prepared by protease digestion for mass spectrometric analysis and PTM containing peptides are enriched for LC-MS/MS. Sample preparation can impact nearly all the later steps in a proteomic study. Therefore, it is critical to design and establish a sample preparation protocol that is robust while reproducible to ensure the scientific integrity of the study. Additionally, a sample preparation workflow with high-throughput capability would be ideal when processing a large set of samples.

The Clinical Proteomics Tumor Analysis Consortium (CPTAC) established and reported a sample preparation protocol for deep-scale MS-based proteomic and phosphoproteomic analysis in 2018^26^. This protocol has been used in various CPTAC-related studies, while the data generated from those studies have been publicly available for the broad scientific community^1,2,6-9^. In the current study, we developed the automation of sample preparation for analyzing proteins and protein modifications protocols, AUTO-SP. We transformed a few essential steps, including protein concentration measurement via BCA assays, protein digestion, and PTM enrichment, in the CPTAC protocol into automated procedures using a liquid handling system. By applying the automation, we could increase sample throughput and reduce potential human errors to enhance the reproducibility and consistency in sample preparation.

In this study, we utilized PDX breast cancer tumor tissues from mouse models (basal-like and luminal subtypes) to demonstrate and evaluate the AUTO-SP. We were able to accurately measure the protein concentration in PDX samples with a coefficient of variation less than 6% among the triplicates. The proteins were further tryptic digested by using AUTO-SP for each PDX sample. Based on the data-independent acquisition (DIA)- MS data, we observed samples of the same breast cancer subtypes were highly correlated (>0.98) with missed cleavage rate between 6 to 7.5%. Additionally, >14,000 ubiquitinated peptides were identified in the PDX samples when using AUTO-SP for ubiquitin enrichment, while unique pathways were enriched from the basal-like and luminal subtypes. Taken together, AUTO-SP demonstrated its efficacy to provide reliable and reproducible sample preparation procedure for MS-based proteomic and PTM analyses.

## METHODS

### Tissue sample

The PDX breast cancer tumor tissues from mouse models, P96 (basal-like) and P97 (luminal), were used in this study. All tumor pieces were cryopulverized, stored at −80 °C until sample preparation for the analysis of global proteomic and protein modifications. The details for preparing bulk cryopulverized PDX tissues can be found in our previous publication^26^.

### Sample processing for protein extraction, tryptic digestion, and ubiquitin enrichment

In this study, BCA analysis, tryptic digestion, and ubiquitin enrichment were carried out by using AUTO-SP and details can be found in the Results. Tissue lysis was performed as previously described^26^. In brief, 400 μL urea lysis buffer (8 M urea, 75 mM NaCl, 50 mM Tris, pH 8.0, 1 mM EDTA, 2 g/mL aprotinin, 10 g/mL leupeptin, 1 mM PMSF, 10 mM NaF, Phosphatase Inhibitor Cocktail 2 and Phosphatase Inhibitor Cocktail 3 [1:100 dilution], and 20 mM PUGNAc) was added to 100 mg of each cryopulverized PDX tissue, followed by repeated vortexing. Lysates were clarified by centrifugation at 20,000 × g for 10 min at 4°C. Lysates from the same breast cancer subtype were pooled together before measuring protein concentrations by BCA assay (Pierce). For the tryptic digestion, pooled samples were aliquoted into a 96-well plate, each well contained 1 mg of protein. In each well, proteins were reduced with 5 mM dithiothreitol (DTT), alkylated with 10 mM iodoacetamide (IAA), diluted 1:3 with 50 mM Tris-HCI (pH 8.0), digested with LysC (Wako Chemicals) at 1 mAU:50 g enzyme-to-substrate ratio and sequencing-grade modified trypsin (Promega) at a 1:50 enzyme-to-substrate ratio. The digested samples were then acidified with 50% formic acid (FA, Sigma) to a pH value of approximately 2.0. Tryptic peptides were desalted on reversed-phase C18 SPE columns (Waters). Aliquoting 1 μg of digested peptides from 12 randomly selected samples to examine the digestion efficiency. All digested samples were dried down in a Speed-Vac. Antibody-based magnetic beads for ubiquitin enrichment came from PTMScan HS Ubiquitin/SUMO Remnant Motif (K-e-GG) Kit (Cell Signaling Technology).

### LC-MS/MS analysis

All the LC-MS/MS data were acquired via EvoSep coupled with timsTOF HT (Bruker) in data-independent acquisition mode. The methods for acquiring global proteomics and ubiquitinated peptides are as follows. A PepSep column of 15 cm x 159 μm (C18, 1.5 μm, Bruker) was used for peptide separation of 30 samples per day at 50°C. The data were acquired under the dia-PASEF mode with a MS1 scan range of 100-1700 m/z, MS2 scan range of 338.6-1338.6 m/z, 1/K0 range of 0.70 to 1.45 V/s/cm^2^, and Ramp time of 85 ms. For the ubiquitinated peptides, we used the same setting as the global peptides, except the MS2 scan range was set between 341.6 to 1216.6 m/z.

### MS data analysis

All the raw files were searched against a UniProt/Swiss-Prot database containing human and mouse proteins (downloaded on 2019/12, 37,405 entries) using directDIA approach in the Spepctronaut (version 18, Biognosys). The search setting is as follows. Mass tolerance of MS and MS/MS was set as dynamic with a correction factor of one. Precursors were filtered by a Q value cutoff of 0.01 (which corresponds to an FDR of 1%). Carbamidomethyl (C) was set as fixed modification. Acetyl (Protein N-term) and Oxidation (M) were set as variable modifications. Variable modifications of GlyGly (K) was set additionally for searching ubiquitinated peptides. The quantity of a peptide was a sum of the quantity of its top 3 precursors, whereas the quantity for a precursor was calculated by summing the area of its top 3 fragment ions at MS/MS level.

Reproducibility among the triplicates was determined based on the Spearman correlation. Specificity of the enrichment of ubiquitination was calculated by summing the abundances of all ubiquitinated peptides and then divided by the total abundances of all identified peptides. Differential analysis was carried out by calculating the median log2 fold changes between two breast cancer subtype groups and two-sided Wilcoxon rank sum was performed with the p-value adjusted via Benjamini-Hochberg method.

## RESULTS

### Overview of AUTO-SP

To increase throughput, minimize human errors, while reduce time and cost for large-scale sample preparation, we have established AUTO-SP, which currently can automate BCA assay analysis, in-solution digestion, and magnetic bead-based enrichment for various protein modifications, including ubiquitination and phosphorylation. **Figure 1** illustrates the general workflow of AUTO-SP for each application. Tumor tissues from the PDX models (P96 and P97) was used to demonstrate the feasibility and reproducibility of the established automated sample preparation platform for the analyses of proteins and protein modifications. For simplicity, we only showed the results of automated ubiquitination enrichment in this study. Of note, we used the Opentrons OT-2 to demonstrate the AUTO-SP in this study, however, we have established Opentrons Flex compatible protocols as well.

**Figure 1.**
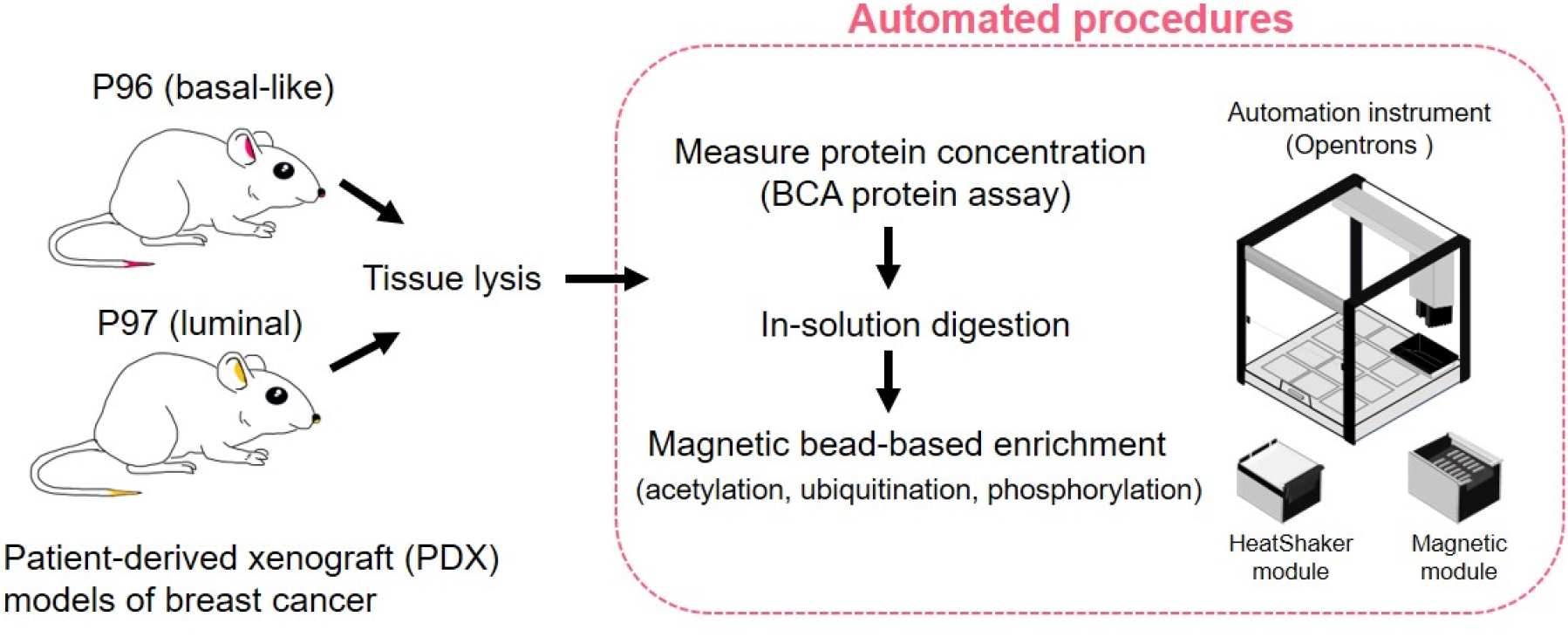
Sample preparation procedures with automated protocols developed.

### Evaluation of automated BCA analysis for protein concentration measurement

Prior to protein digestion, measuring protein concentration in a sample is essential to ensure adding an adequate amount of protease (e.g., trypsin) to avoid incomplete or over digestion of proteins. Using BCA protein assay to quantify protein amount is a widely-adopted approach in the proteomic studies. BCA reagent is added and incubated with diluted samples before reading protein concentration in each sample. Triplicate of each sample is usually used for ensuring accuracy of measurement. The whole process is simple and easy to operate when handling only a small set of samples. However, large-scale proteomic studies could have more than 100 samples that need to be analyzed. Manually processing BCA assays will then become time-consuming and human errors are likely to occur when transferring samples from individual vials to a 96-well plate. Therefore, we established the automated BCA assay protocol on the Opentrons platform to increase throughput, while reduce human and random errors.

After urea-based tissue lysis, 2 uL from pooled P96 and pooled P97 were diluted using HPLC water (1:20 ratio) and placed into a 96-well plate (i.e., source plate) along with the BSA protein standards (**Figure 2A**). **Figure 2B** demonstrates the layout on the Opentrons for this particular protocol. The BCA protocol can be started when all required labware are placed in the designated slots. As shown in **Figure 2C**, the measured protein concentration was consistent between pooled P96 samples as well as among pooled P97 samples. Moreover, majority of the samples with coefficient of variation (CV) less than 3% among the triplicates were observed, while the maximum CV was 5.08% (**Figure 2D**), further indicating the successful execution of the automated BCA analysis.

**Figure 2.**
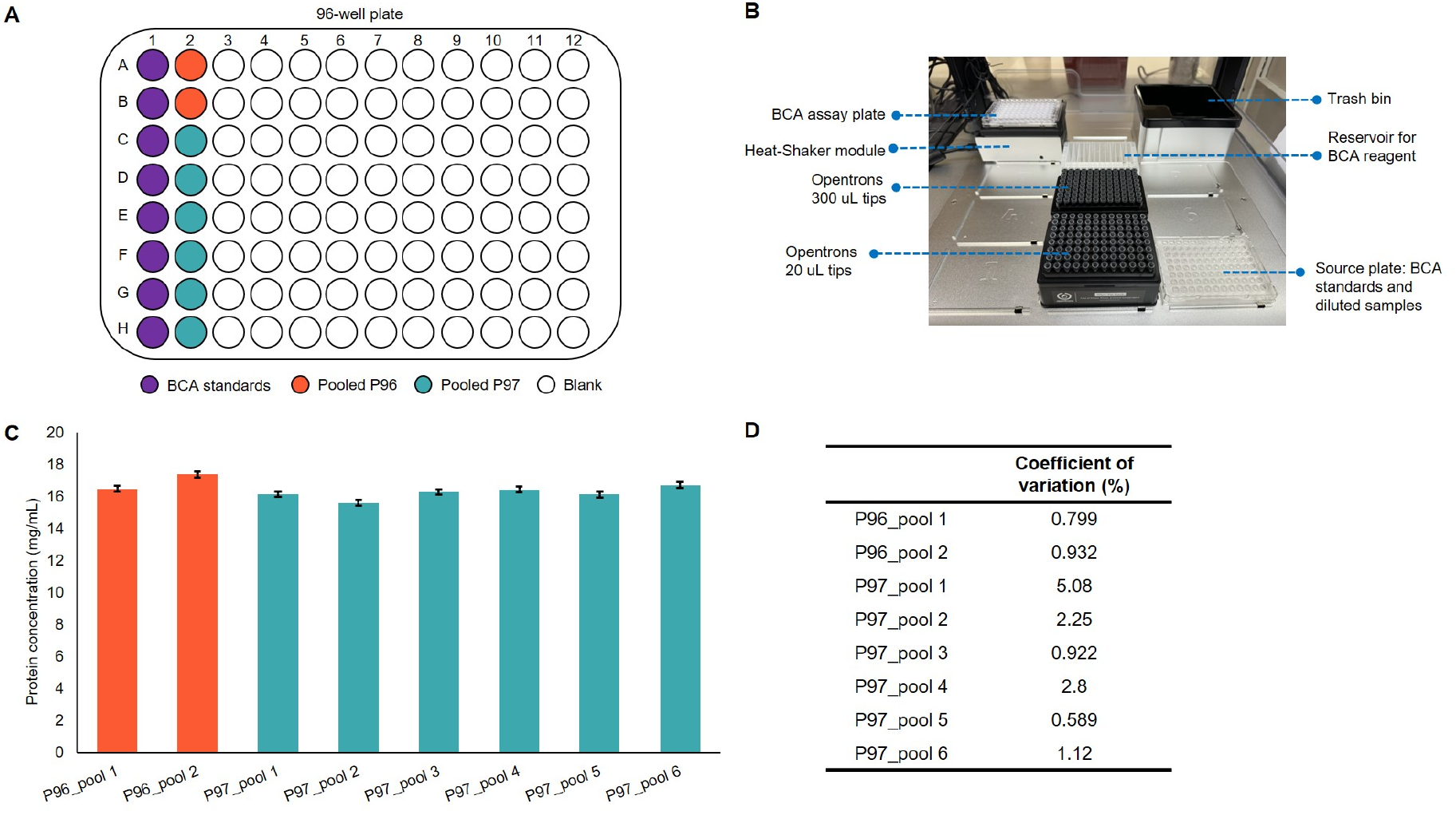
BCA analysis of pooled samples. **A.** Layout on a 96-well plate for holding BCA standards and diluted samples for the current study. **B**. Layout on the Opentrons used in the current study. **C**. Protein concentration measured in each pool sample. **D**. Coefficient of variation among the triplicates of each pool sample.

### Assessment of in-solution digestion

For the bottom-up proteomics, proteins and protein modifications are identified and quantified based on the peptides derived from each protein by enzymatic digestion. The amount of a protease to be added is estimated according to the BCA analysis as described above. In this study, we used trypsin for protein digestion, however, the protocol can be adapted to protein digestion using other proteases.

The pooled P96 and P97 were distributed to a 96 deep-well plate (source plate) that each well contained approximately 1 mg of proteins (**Figure S1A**). The layout of the labware on the Opentrons was similar to the BCA analysis but the source plate was placed on the Heat-Shaker module instead. All the reagents, buffer, and proteases were added automatically to the source plate on the Opentrons. The incubation of IAA (45 min), LysC (2h), and Trypsin (overnight, ∼16h) were carried out on the Heat-Shaker at room temperature at a speed of 500 rpm, except for the DTT. The Heat-Shaker from the Opentrons can provide heat to the bottom of a plate. However, to ensure temperature uniformity during the incubation of DTT, the source plate was moved from the Opentrons to an incubator shaker for 1h incubation at 37°C. Of note, during transferring and incubation of IAA, the Opentrons was completely covered to avoid light interfering in alkylation of proteins.

We randomly selected 12 wells to evaluate the efficiency of AUTO-SP on in-solution digestion (**Figure S1A**). High reproducibility was observed among the wells containing the same sample types since the Spearman correlation was ≥0.98 (**Figures 3A, S1B**) with median coefficient of variation below 10% for P96 and P97 (**Figure 3B, S1C**) indicating that AUTO-SP could provide consistent results. Moreover, the missed cleavage rate was between 6.1 to 7.5%, regardless of the sample types (**Figure 3C**). The total number of peptides and proteins identified from P96 and P97 samples are as shown in **Figure 3D**. On average, 106,700 and 11,700 peptides and protein groups were identified across P96 and P97.

**Figure 3.**
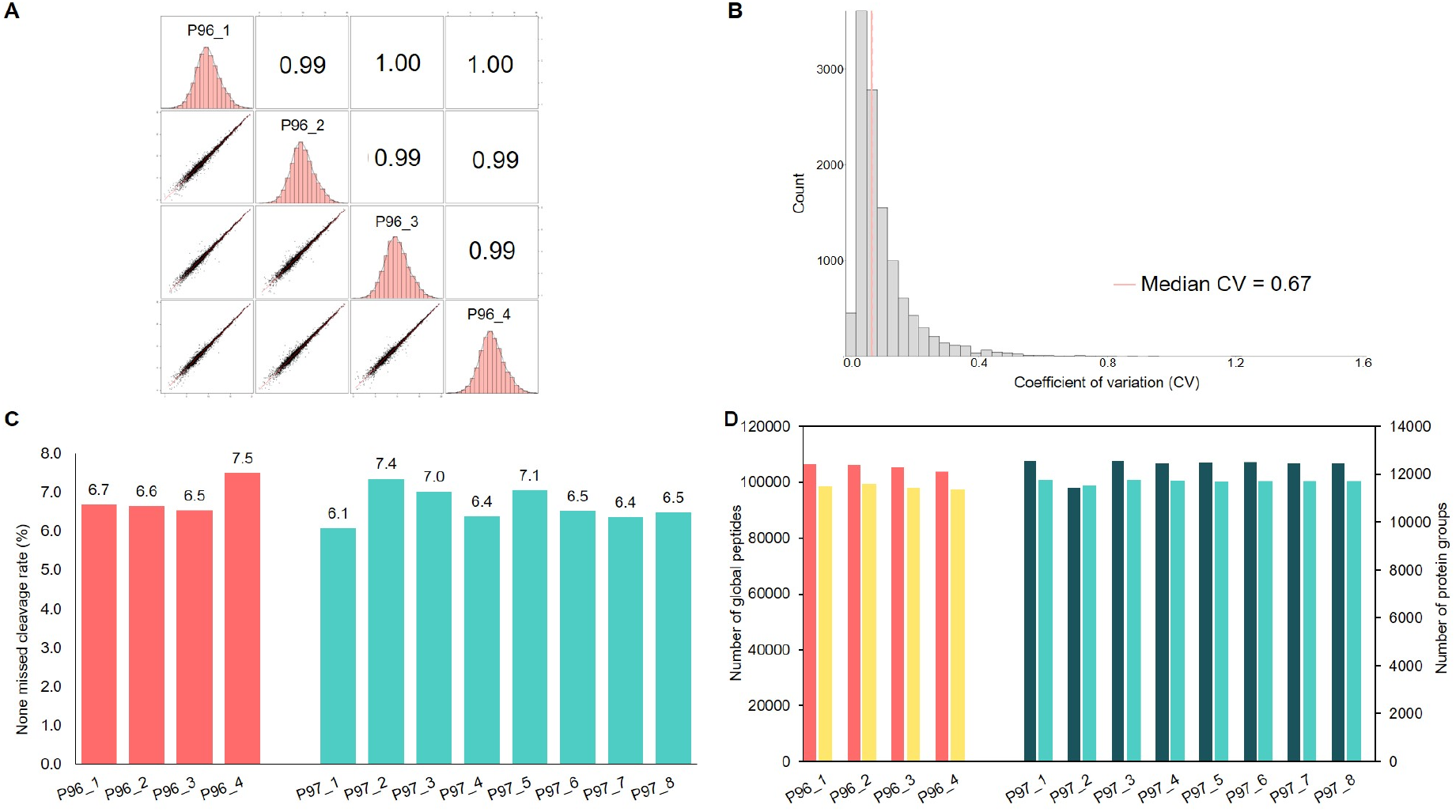
In-solution trypsin digestion using AUTO-SP. **A.** Reproducibility of protein digestion on AUTO-SP for basal subtype. **B**. Coefficient of variation of basal subtype. **C**. None missed cleavage rate of each sample. **D**. Total identified global peptides and protein groups for each sample.

Overall, AUTO-SP produced consistent and reliable results, while supported high throughput of protein digestion.

### Magnetic bead-based ubiquitin antibody enrichment analysis

In addition to BCA analysis and in-solution digestion, AUTO-SP also facilitates enrichment of protein modifications via magnetic beads, accommodating both antibody-based and non-antibody-based beads. In the current study, we demonstrated the efficacy of protein modification enrichment on AUTO-SP using antibody-based magnetic beads to isolate ubiquitinated peptides from P96 and P97.

A total of 30 P96 and 61 P97 global peptide samples were generated using the AUTO-SP as described above. We randomly selected 9 from each breast cancer subtype and made pooled P96 and pooled P97 for the enrichment. For each subtype, 6 replicates were prepared from the pooled samples for each subtype. Each replicate started with approximately 300 μg of peptides and 5 μL magnetic bead slurry. The enrichment process on AUTO-SP included the following: (1) transferring samples to a 96-well plate on the Magnetic Module of Opentrons, (2) incubating samples with the beads, (3) taking flow-through, (4) washing beads with IAP wash buffer (2 cycles), (5) washing beads with PBS (2 cycles), and (6) eluting ubiquitinated peptides from the beads (100 μL 0.15% TFA twice with final total volume of 200 uL). Pipette mixing was used to thoroughly mix the samples with antibody beads during incubation and elution as well as during beads washing.

By using the magnetic bead-enrichment protocol of AUTO-SP, a total of 16,531 non-redundant ubiquitinated peptides were identified from 6 replicates of P96, with an averaged identification of 14,770 ubiquitinated peptides (**Figure 4A**). The enrichment specificity was ranged from 19% to 34%, with a median at 24.8% (**Figure 4B**). Similarly, 16,701 ubiquitinated peptides were identified across the replicates of P97 with an enrichment specificity ranging from 23.7% to 33.7% (median = 28.7%).

**Figure 4.**
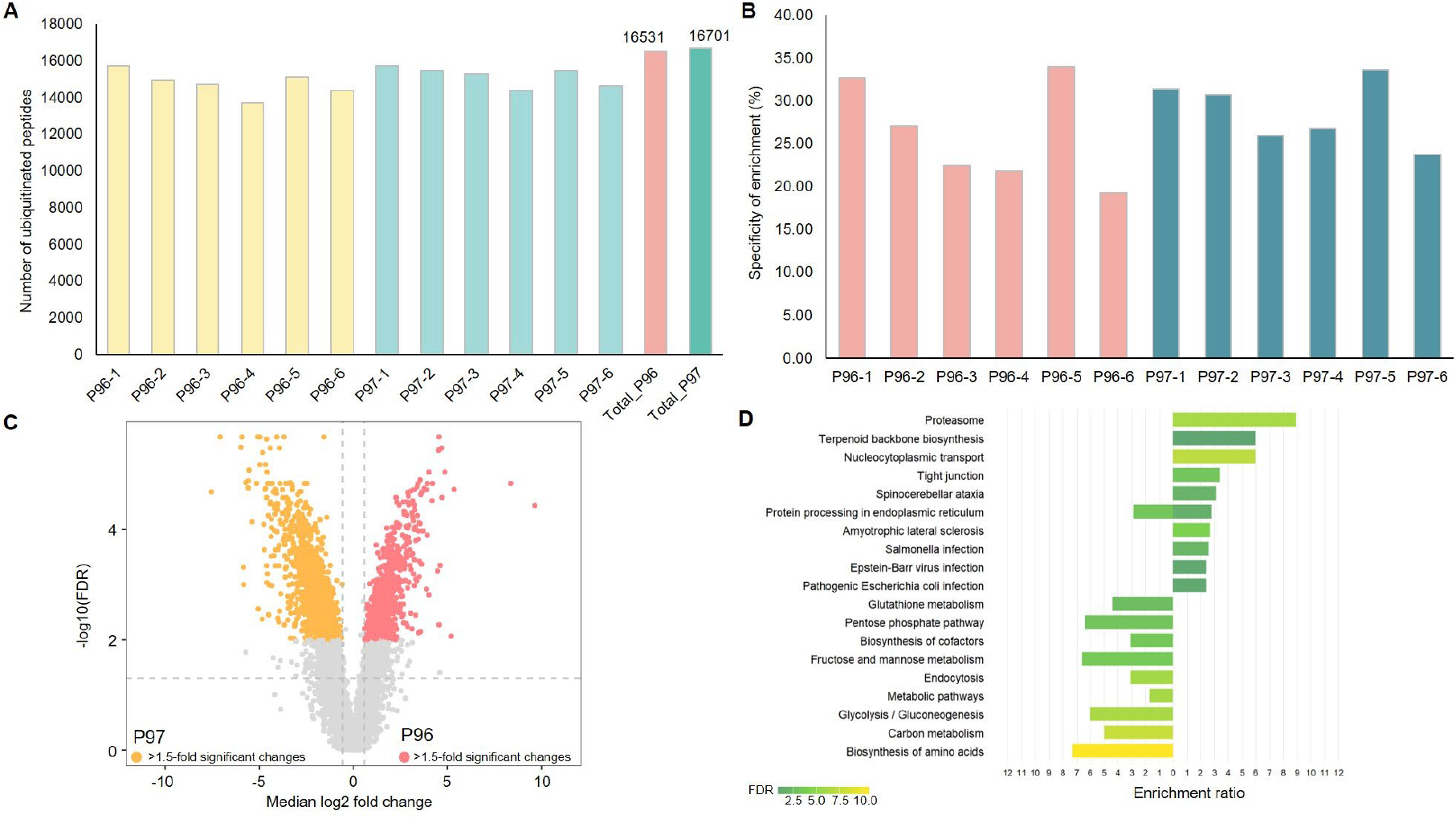
Ubiquitin enrichment via AUTO-SP. **A.** Number of identified ubiquitinated peptides in each sample and total non-redundant ubiquitinated peptides identified across P96 and P97. **B**. Ubiquitin enrichment specificity. **C**. Comparison of P96 and P97 based on the enriched ubiquitinated peptides. **D**. Enriched KEGG pathways based on unique proteins with ubiquitinated peptides that were differentially expressed in P96 and P97.

Ubiquitination of proteins regulates various cellular processes, including proteasomal degradation, intracellular trafficking, and DNA repair^27,28^. An association between cancer development and dysreuglation of protein ubiquitination has been reported previously^27,29,30^. Comparing the two PDX breast cancer subtypes, 719 ubiquitinated peptides (originated from 421 proteins) with elevated expression in P96 relative to P97. On the other hand, 1293 ubiquitinated peptides (originated from 573 proteins) with higher abundance in P97 than P96 (**Figure 4C**). From the differentially expressed ubiquitinated peptides, we found that unique pathways were enriched for P96 and P97 (**Figure 4D**).

The aforementioned results demonstrated the efficacy of using AUTO-SP for PTM enrichment and its capability to handle large-scale samples.

## DISCUSSION

Sample preparation is one of the critical components to the success of a study. To generate consistent and reproducible data while support high sample throughput, we established the AUTO-SP, which adapted some of the essential steps in the CPTAC protocol published in 2018 by converting them into automated procedures for deep-scale MS-based quantitative proteomic and PTM analyses.

We used the PDX tumor tissues of P96 and P97 to evaluate the performance of AUTO-SP on protein concentration measurement, protein tryptic digestion, and ubiquitinated enrichment using magnetic antibody beads. We found that AUTO-SP produced highly reproducible and consistent results among the samples from the same subtypes, while it was capable of simultaneously handling > 80 samples. We identified more than 10,000 proteins while replicates showed high correlation (>0.98), which was comparable to the 2018 CPTAC protocol^26^. Moreover, there were notable significant expression differences of the ubiquitinated peptides between the two subtypes. Using the proteins of differentially expressed ubiquitinated peptides, different pathways were enriched. For example, nucleocytoplasmic transport pathway was emerged from P96, while carbon metabolism pathway was enriched in P97. Although further examination of the biological significance of these pathways would be required, however, these results indicated the successful application of AUTO-SP for ubiquitinated peptide analysis and its potential for analyzing large-scale clinical samples.

In conclusion, AUTO-SP could support high-throughput and reliable automated sample preparation for MS-based proteomic and protein modification analyses of clinical samples.

## ACKNOWLEDGEMENTS

This work was supported by the National Institutes of Health, National Cancer Institute, Clinical Proteomic Tumor Analysis Consortium (CPTAC, U24CA271079), Early Detection Research Network (EDRN, U2CCA271895), and Pancreatic Cancer Detection Consortium (PCDC, U01CA274514).

