## Supplementary material for "AUTO-SP: automated sample preparation for analyzing proteins and protein modifications": Figure S1

**Supplementary file 1**

### AUTO-SP: automated sample preparation for analyzing proteins and protein modifications

T. Mamie Lih[1,*], Liyuan Jiao[1], Lijun Chen[1], Jongmin Woo[1], Yuefan Wang[1], Hui Zhang[1,2,3]

1 Department of Pathology, Johns Hopkins University School of Medicine, Baltimore, Maryland, United States.

2 Department of Oncology, Sidney Kimmel Cancer Center at Johns Hopkins Medical Institutions, Baltimore, Maryland, United States.

3 Department of Urology, Johns Hopkins University School of Medicine, Baltimore, Maryland, United States.

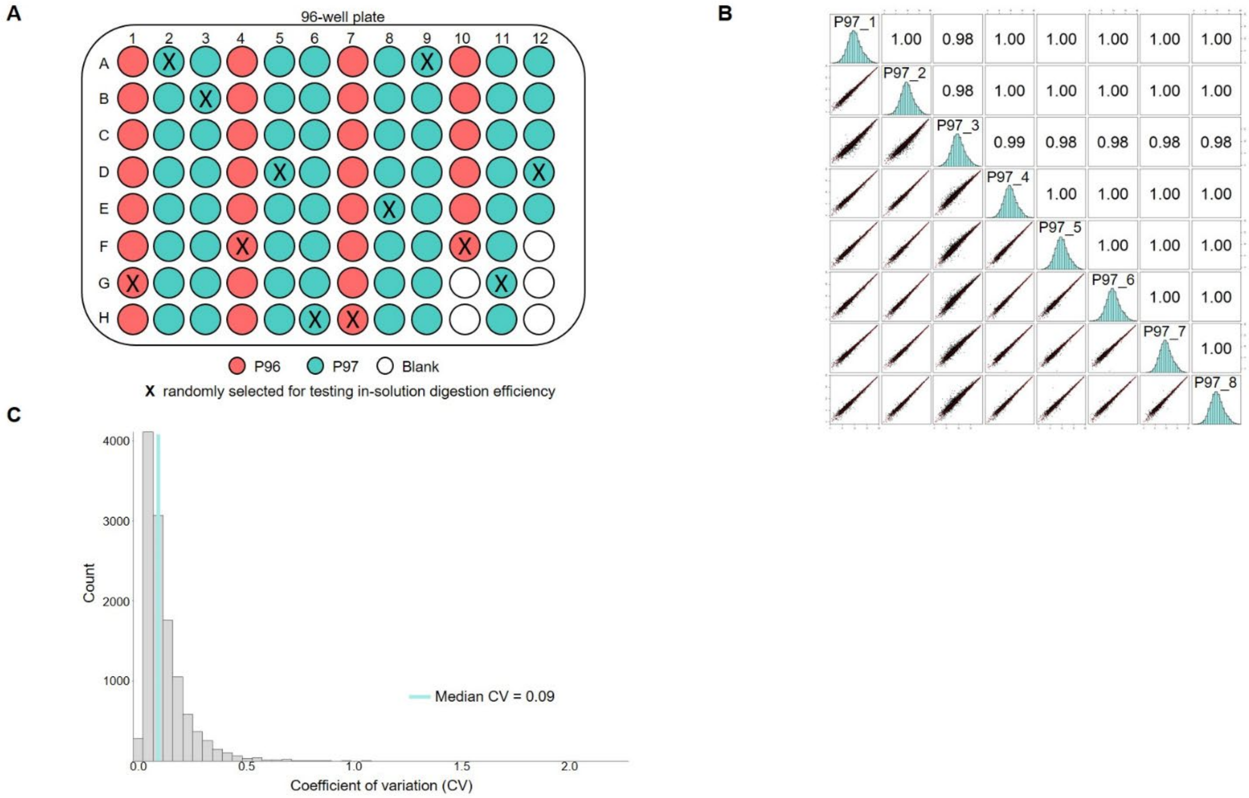


**Figure S1.** Protein digestion using trypsin on AUTO-SP related to Figure 3. **A.** Sample layout on a 96-well plate on the AUTO-SP for protein digestion. **B.** Reproducibility of protein digestion on AUTO-SP for P97. **C.** Coefficient of variation of P97 samples.
